# Correlation between the aberrant human testicular germ-cell gene expression and disruption of spermatogenesis leading to male infertility

**DOI:** 10.1101/394049

**Authors:** Arka Baksi, Ruchi Jain, Ravi Manjithaya, S S Vasan, Paturu Kondaiah, Rajan R. Dighe

**Affiliations:** Department of Molecular Reproduction, Development and Genetics, Indian Institute of Science, Bangalore, India; Molecular Biology and Genetics Unit, Jawaharlal Nehru Centre for Advanced Scientific Research, Bangalore, India; Manipal Fertility, Bangalore, India

## Abstract

Spermatogenesis is characterized by sequential gene-expression at precise stages in progression of differentiation of the germ cells. Any alteration in expression of the critical genes is responsible for arrest of spermatogenesis associated with infertility. Inspite of advances the differential gene expression accompanying spermatogenesis, the corresponding regulatory mechanisms and their correlation to human infertility have not been clearly established. This study aims to identify the gene expression pattern of the human testicular germ cells from the patients either with obstructive azoospermia with complete intra-testicular spermatogenesis or non-obstructive azoospermia with spermatogenesis arrested at different stages and correlate the same to infertility. The testicular transcriptomes of 3 OA and 8 NOA patients and pooled testicular RNA (commercial source) were analyzed for their differential gene expression to identify potential regulators of spermatogenesis and the results were further validated in all of the 44 patients clinically diagnosed with azoospermia undergoing sperm retrieval surgery over the study period and 4 control samples included in this study. Analyses of the differential transcriptome led to identification of genes enriched in a specific testicular cell type and subsequently, several regulators of the diploid-double-diploid-haploid transitions in the human spermatogenesis were identified. Perturbations in the expression of these genes were identified as the potential causes of the spermatogenic arrest seen in azoospermia and thus the potential mediators of human male infertility. Another interesting observation was the increased autophagy in the testes of patients with non-obstructive azoospermia. The present study suggests that the regulation of the diploid-double-diploid-haploid transition is multigenic with the tandem alteration of several genes resulting in infertility. In conclusion, this study identified some of the genetic regulators controlling spermatogenesis using comparative transcriptome analyses of testicular tissues from azoospremic individuals and showed how alterations in several genes results in disruption of spermatogenesis and subsequent infertility. This study also provides interesting insights into the gene expression patterns of the Indian population that were not available earlier.

## Introduction

Spermatogenesis is an extraordinary process of cell differentiation in which the testicular germ cells undergo series of mitotic divisions, followed by a meiotic division and transformation of the haploid cells into spermatozoa which eventually attain motility. Regulation of spermatogenesis occurs at different levels, the first being the extrinsic level wherein the gonadotropins and testosterone regulate gene expression in the germ cells sustaining their survival and progression of differentiation ^1^ and blockades in their action results in apoptotic death of the germ cells ^2-4^. The second level is the interactive regulation that involves communications between the somatic cells such as the Sertoli cells and the germ cells. The third level of regulation is the intrinsic gene expression associated with the germ cell differentiation ^1,5^. These differentiation events are characterized by sequential gene-expression wherein the specific genes are turned ‘on’ or ‘off’ at precise stages facilitating the progression of differentiation of the germ cells. Any alteration in the expression of the genes appears to be responsible for arrest of spermatogenesis associated with infertility ^6-12^. However, the critical genes required for progression of spermatogenesis and the genes whose altered expression leads to its arrest have not been clearly identified ^13^. Identification of such genes is essential for understanding the molecular defects in arrested spermatogenesis and possible treatment of the consequent infertility. However, such a task has proved to be difficult due paucity of the human testicular tissue from the normal, fertile individuals, as well as, the infertile males with arrested spermatogenesis. The other contributing factor is the complex organization of spermatogenesis and the unique environment present in the seminiferous tubules that has not been replicated *in vitro*. It is also not possible to culture the germ cells and make them undergo differentiation *in vitro.*

The infertile patients with the spermatogenesis arrested at any stage of differentiation provide an interesting paradigm for investigating alterations in the testicular transcriptome that could be correlated to spermatogenic arrest. In the present study, we investigated gene-expression changes in the testes of the infertile patients with non-obstructive azoospermia having spermatogenic arrest either at the diploid or double-diploid stages and compared it to the germ cell gene expression patterns of the obstructive azoospermic (OA) patients who exhibited complete spermatogenesis. This allowed identification of genes which could potentially be responsible for the arrest. The study demonstrates that alterations in the expression of several regulators of spermatogenic progression are the potential causes of germ-cell arrest leading to infertility.

## Material and Methods

### Patient Samples

The testicular biopsies of infertile patients were obtained from Manipal Ankur Fertility Clinic, Bangalore with appropriate clearances from the Institutional Review Board of Manipal Ankur Fertility Clinic and the Human Ethics Committee of the Indian Institute of Science, Bangalore. All experiments were performed in accordance with the relevant guidelines and regulations with prior approval.

### RNA isolation, RT-PCR and real-time PCR

The total RNA from the testicular tissue was isolated using TRI reagent (Sigma-Aldrich, USA) as per the manufacturer’s protocol. cDNA was generated from 2µg of RNA using the Revert Aid First Strand cDNA Synthesis kit (Thermo Fisher Scientific, USA). qRT-PCR was performed using the ABI Prism 7900HT sequence detection system. cDNA equivalent to 20ng RNA was used for qRT-PCR analysis using Dynamo SYBER green 2X mix (Finnzymes, Finland) along with gene-specific primer pairs (Table S1). RPL35 expression was used for normalization. Differential expression of genes was determined using the following formula:

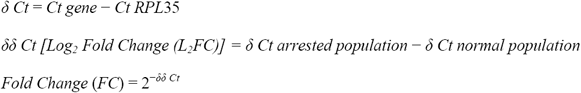

### Transcriptome Profiling using Microarray

The quality and quantity of RNA was analyzed using Qubit RNA Assay Kit in Qubit flurometer (Thermo Fisher Scientific, USA) followed by Agilent 2100 Bioanalyzer (Agilent Technologies, USA). Samples with RNA-integrity values>6 were chosen for microarray and biotin-labeled complementary-RNA (c-RNA) was synthesized from each using the TargetAmp-Nano Labeling Kit for Illumina Expression BeadChip (Illumina Inc. USA). The cRNA was hybridized onto an Illumina HumanHT-12v4 chip (Illumina Inc. USA) and signal detection was performed by incubating with Cy3-streptavidin conjugate in HiScanSQ System (Illumina Inc. USA). The total human testicular RNA pooled from 5 individuals of Asian origin (Clonetech, USA) was used as the control. The preprocessing and identification of differentially expressed genes are provided in the supplementary section.

### Data Availability

The complete microarray data are available at GEO database, accession No GSE10886

### Gene-Enrichment analysis

The enrichment of the population specific gene sets to GO terms was analyzed using the ‘Enrichr’ tool (http://amp.pharm.mssm.edu/Enrichr/) ^14^. Enrichment of biological processes was tested for the haploid, diploid and the double-diploid-enriched genes. Pathway analysis was performed by uploading to KEGG (http://www.genome.jp/kegg/) ^15^.

### Transcription factor Binding site analysis and Network construction

The testicular cell population-enriched genes were searched for the putative transcription factor binding sites using Mapper2^16^. Network maps were created through String database^17^ for each population-enriched gene set (diploid, double-diploid and haploid) and their respective differentially-expressed transcription factors (only the putative transcription factors that were differentially expressed in the microarray analysis were selected).

### Immunohistochemistry and TUNEL assay

Immunohistochemistry was performed as described in Pant *et al* ^18^. The primary antibodies used were Anti-EGR2 (abcam), Anti-RFX2 (Sigma-Aldrich), Anti-CDKN1A/p21 (Cell Signaling Technology) and Anti-LC3B (Sigma-Aldrich). In-situ Apoptosis Detection Kit (Abcam) was used to check for the apoptotic cells in the tissue sections according to the Manufacturer’s protocol. Additional details are provided in the supplementary section.

### Image Capture

All images were captured using the Zeiss microscope and processed by Zeiss Axiocam 4.3 software.

### Statistical Analysis

Unpaired t-test with Welch’s correction was performed to assess statistical significance for the expression of the selected genes in qRT-PCR. GraphPad Prism V software was used to analyze data for statistical significance. p value ≤ 0.01, 0.001, 0.0001 are represented as ^*, **, ***^ respectively.

## Results

### Patients

The classification the infertile patients analyzed in this study has been described in details in Baksi et al^19^. Forty four infertile patients were first classified into OA or NOA based on clinical inputs and further classified into three distinct groups based their germ cell flow cytometry coupled with marker gene analysis as shown in the previous publication from the laboratory. In addition, 4 patients, previously known to be fertile, but undergoing surgical procedures unrelated to infertility were considered as the controls. Six patients exhibited only diploid cells indicating diploid/pre-meiotic arrest (Group III or Group D), while 24 patients showed presence of the diploid and double-diploid cells indicating double-diploid/meiotic arrest (Group II or Group DT). Fourteen patients with OA showed presence of the diploid, double-diploid and haploid cells (Group I or Group DTH) indicating complete spermatogenesis. The control, fertile individuals also exhibited full spermatogenesis with all three germ cell populations^19^.

### Transcriptome profiling

The total RNA samples from Groups D (3), DT (5) and DTH (3) and the total testicular RNA (N) (pooled from 8 individuals, Clontech, USA) (Figure S1) were subjected to transcriptome analysis by Microarray and the genes expressed differentially between the groups were identified (Table S2). Seventeen genes representing the highest differential-expression and the testicular cell-specific markers were validated by qRT-PCR (Figure S2). The gene-expression profiles were further analyzed to identify the testicular cell-specific enriched genes (Figure S3 and S4).

### Gene set enrichment and pathway analysis

The population-enriched gene sets were analyzed for enrichment to GO terms. The diploid-enriched genes showed predominance for early developmental processes active in the spermatogonia ^3,4,20,21^. The double-diploid-enriched genes were enriched for the biological processes such as inflammation, stress response and movement of cellular components. The haploid-enriched genes showed enrichment for the male-gamete formation, spermatid development, ciliary movement and sperm motility (Table S3). Pathway analysis through KEGG also provided the pathways regulated by the population-enriched genes (Table S4). These results indicated that the annotation of the cell population-enriched gene sets was in accordance with their function.

### Transcription Factors and Network Maps for population-enriched genes

Network maps were constructed using the population-enriched genes and their TFs (Figures S5-7). The genes having ≥4 connections were identified from these networks of which 15, 13 and 58 genes were unique to the diploid, double-diploid and haploid subsets respectively, while 12 genes were common to all three groups (Figure S8).

### Identification of the population-enriched genes possibly responsible for arrest of spermatogenesis and validation at the transcript levels

The diploid-enriched genes were mainly the regulators of cell-cycle and proliferation, while the double-diploid and haploid-enriched genes were potential regulators of the double-diploid to haploid transition ^22-33^ (Figure S9A-C). As these observed alterations could potentially cause the spermatogenic arrest, these genes have been referred to as ‘crucial’ population-enriched genes. Subsequently, these crucial genes were validated at the transcript level in all 4 control and 44 patients’ samples and their expression patterns corroborated the microarray analysis (Figure 1A-C).

**Figure 1.**
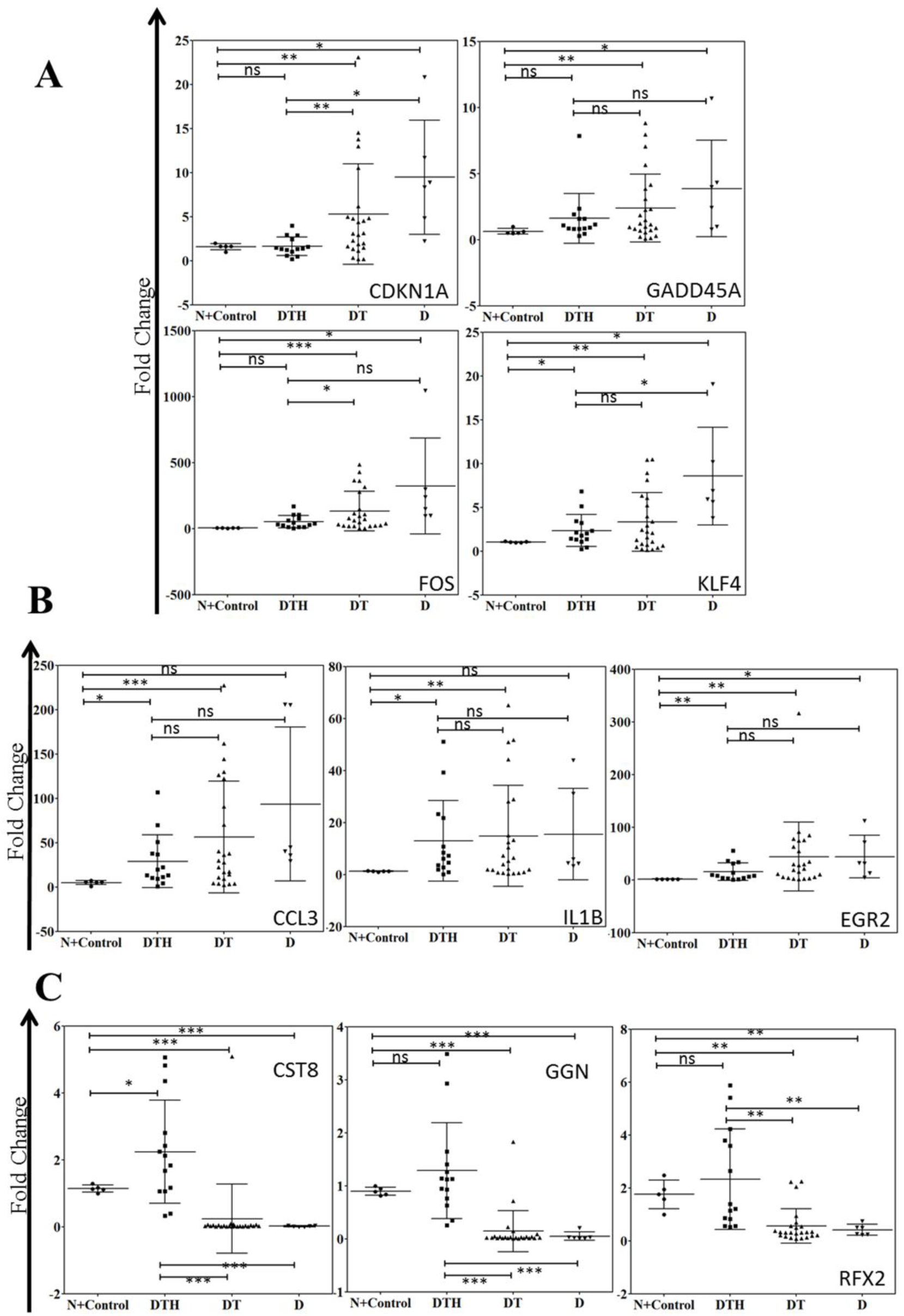
Transcript levels of testicular cell population-enriched genes crucial for diploid to double-diploid to haploid transition across all patient samples. The transcript levels of the crucial testicular cell population-enriched genes were ascertained in all the 44 patient samples (samples included or not-included in the microarray) and the control using specific primers in qRT-PCR and the significance was calculated using the unpaired t-test with Welch’s correction. **A. Transcript levels of diploid-enriched genes crucial for diploid to double-diploid transition. B. Transcript levels of double-diploid-enriched genes crucial for double-diploid to haploid transition. C. Transcript levels of haploid-enriched genes crucial for double-diploid to haploid transition.** N: Pooled RNA control, Control: patients with proven fertility, DTH: obstructive azoospermia patients with complete intra-testicular spermatogenesis, DT: meiotic arrest patients with diploid and double-diploid cells in their testes, D: pre-meiotic arrest patients with only diploid cells in their testes; P > 0.05- ns; P ≤ 0.05-*; P ≤ 0.01-**; P ≤ 0.001-***

### Validation of selected genes using immunohistochemistry

The hubs of the diploid and double-diploid-enriched altered genes, CDKN1A (Figure S9A) and EGR2 (Figure S9B), and RFX2 (Figure S9C), whose aberration could have wide-ranging effects on the cell-cycle were validated at the protein level using specific antibodies. Staining of CDKN1A was seen only in Group D, whereas Groups DT and DHT exhibited poor staining (Figure 2A). The nuclear staining for EGR2 was seen in the Group DT with no staining in Group DTH and only a faint signal seen in Group D (Figure 2B). RFX2 staining was observed in the double-diploid and haploid cells of Group DTH, while no expression was seen in Group DT spermatocytes. Group D sections also did not stain for RFX2 due to absence of the double-diploid cells (Figure 2C). Thus, the differences seen at the transcript level were further confirmed at the protein level.

**Figure 2.**
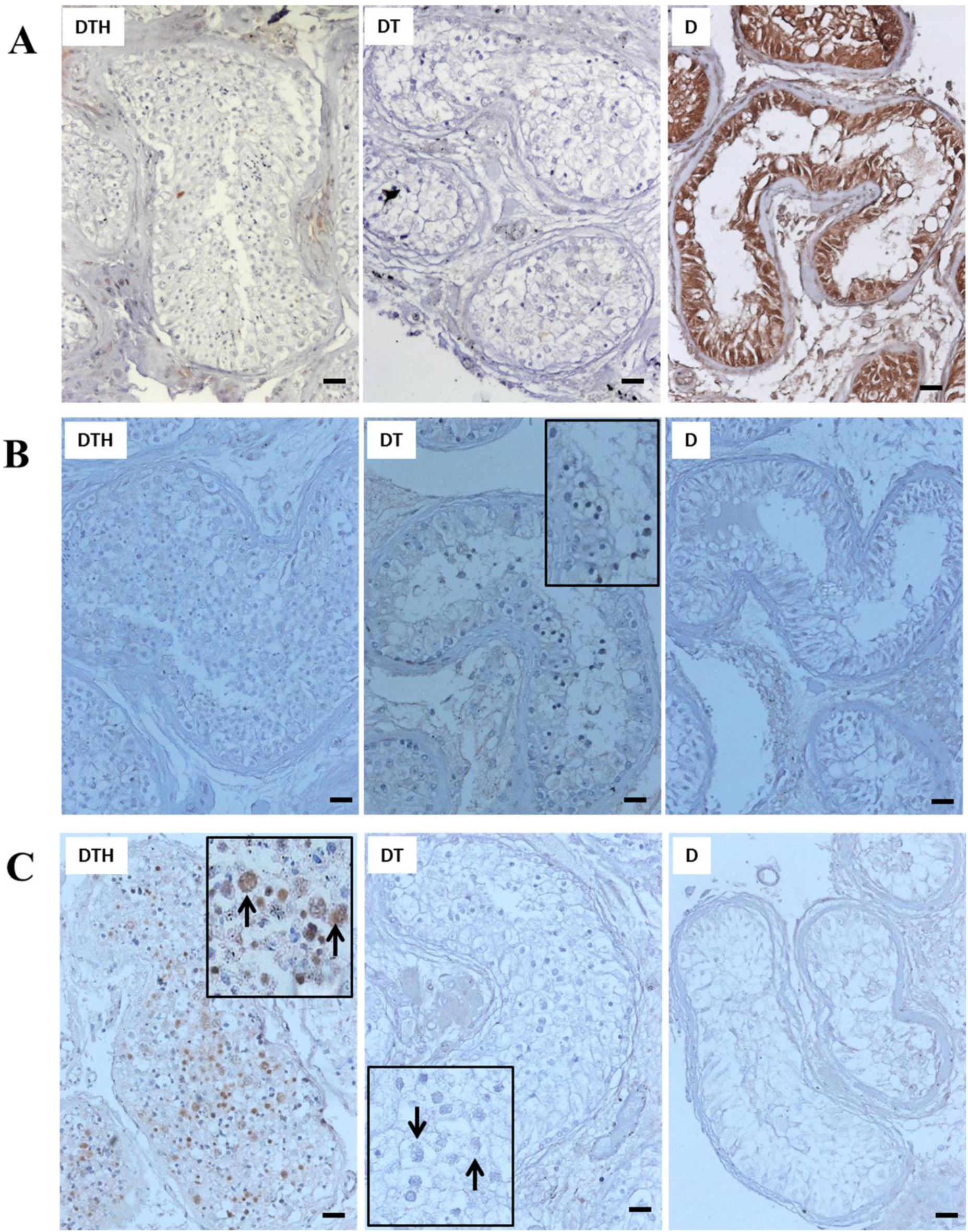
Representative photomicrographs of expression of A. CDKN1A, B. EGR2 and C. RFX2 proteins in testicular tissue sections of Groups DTH, DT and D patients. Expression of CDKN1A, EGR2 and RFX2 was ascertained at the protein level by immuno-histochemistry. **B. Inset**: magnified image showing nuclear staining of EGR2 in the Group DT sample. **C. Inset**: magnified image [200X] showing differences in expression of RFX2 in the spermatocytes present in Group DTH and DT patients. The spermatocytes or meiotic cells are marked with black arrows. DTH: obstructive azoospermia patients with complete intra-testicular spermatogenesis, DT: meiotic arrest patients with diploid and double-diploid cells in their testes, D: pre-meiotic arrest patients with only diploid cells in their testes; Scale: 50µm, Magnification: 100X.

### Status of apoptosis and autophagy in the azoospermic patients

The status of apoptosis and autophagy was ascertained in the patient samples as possible clearance mechanisms for the arrested germ-cells. No appreciable difference was seen in the status of apoptosis in the patients belonging to all the groups (Table-S5 and Figure 3A). For autophagy, no difference was observed at the transcript level of the autophagy-related genes (Table S6). However, a significant increase in LC3B staining was observed in the Groups D and DT while the Group DTH showed a faint staining (Figure 3B).

**Figure 3.**
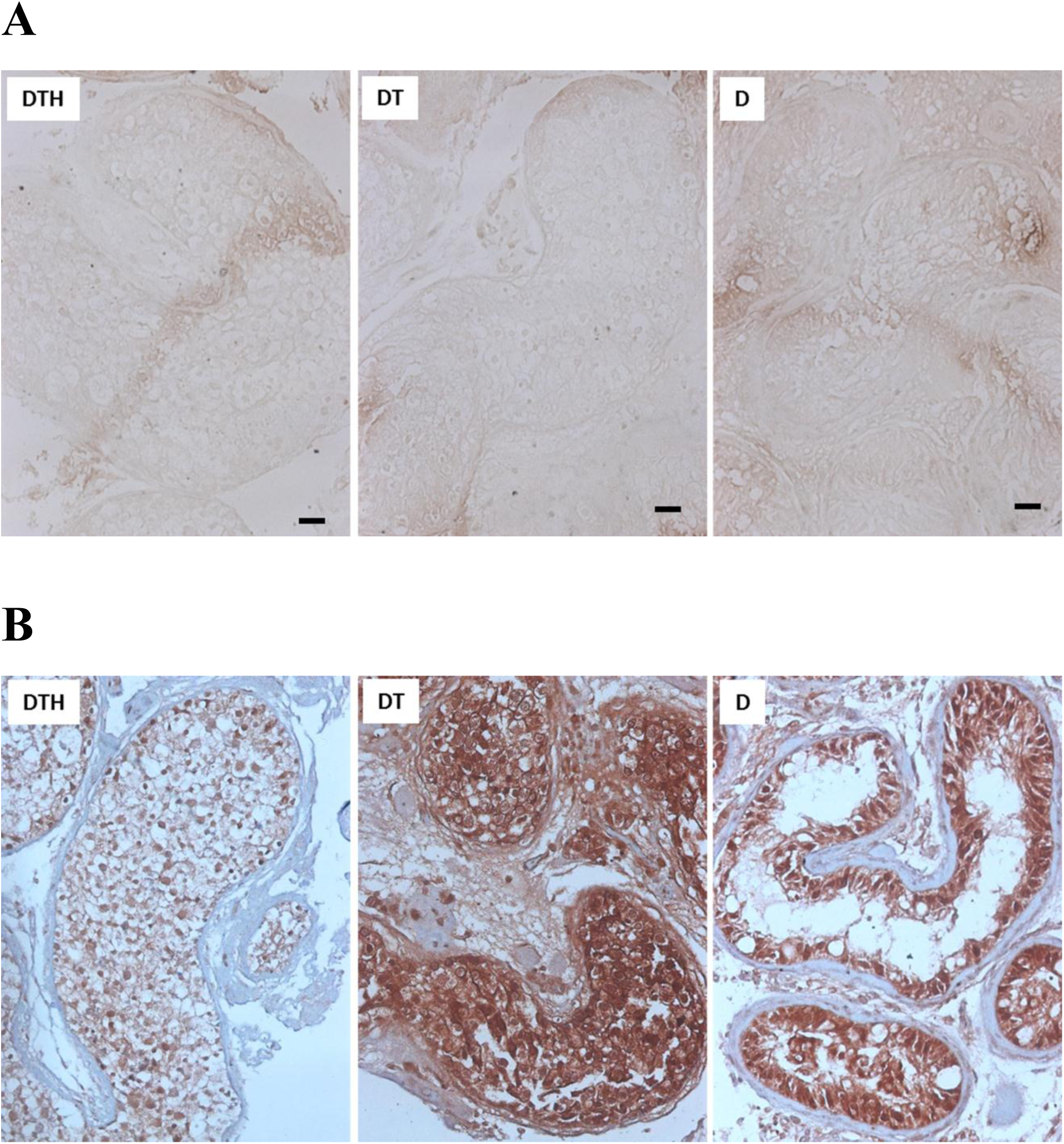
Representative photomicrographs showing the status of A. Apoptosis and B. Autophagy in testicular tissue sections of patients from Groups DTH, DT and D. A. The status of apoptosis was ascertained by TUNEL assay. **B.** The status of autophagy was determined in the different groups of patients by immuno-histochemical staining for LC3B. DTH: obstructive azoospermia patients with complete intra-testicular spermatogenesis, DT: meiotic arrest patients with diploid and double-diploid cells in their testes, D: pre-meiotic arrest patients with only diploid cells in their testes; Scale: 50µm, Magnification: 100X.

## Discussion

Arrest of spermatogenesis constitutes approximately 10% of infertile male patients and devising novel therapeutic strategies to treat such infertility has been an extremely challenging task. Lack of understanding of the molecular mechanisms that govern the spermatogenic differentiation and inability to manipulate germ cell differentiation *in vitro* have been the major impediments in developing such strategies. The testicular tissues from the infertile patients with spermatogenesis arrested at different stages of differentiation provide an interesting model for understanding for understanding the molecular mechanisms operative in the human testis. In this study, an extensive analysis of testicular gene expression patterns of the NOA patients led to identification of some of the critical genes that play roles in progression of spermatogenesis and aberrant expression of these genes may be responsible for arrest of germ cell differentiation. Presented below is the discussion on some of the genes with aberrant expression which could be responsible for arrest of spermatogenesis based on the information available in the literature with both human and rodent models.

CDKN1A, an inhibitor of cyclins ^34^, exhibited higher transcript levels in all Group D patients and formed the hub of the network of the diploid-enriched genes (Figure S9A). Over-expression of CDKN1A is associated with inhibition of cell-proliferation and the cell-cycle arrest (mediated by nuclear CDKN1A) ^22,35^. In mice, CDKN1A maintains the spermatogonial stem cell proliferation as its overexpression inhibits germ-line stem cell proliferation ^36^. The CDKN1A-null mice did not undergo G1-arrest in response to induced DNA damage^37,38^ identifying it as a mediator of the G1 check-point. Interestingly, CDKN1A deficiency does not affect the stem cell number or self-renewal mechanism^36^, but the undifferentiated spermatogonia exhibit up-regulation of CDKN1A upon DNA damage^39^. The CDKN1A knockout mice showed 35% increase in their testicular weight due to increased proliferation of the germ cells. In the ATM-deficient mice spermatogenesis is halted at the diploid stage via CDKN1A mediated inhibition of germ-cell proliferation^39^. In human testis, CDKN1A levels were virtually undetectable in the germ cells^40^ while the non-dividing Leydig cells showed a moderate expression^41^. In the present study, all 6 Group D patients showed an elevated expression of CDKN1A compared to DTH and N, both at the transcript (Figure 1A) and protein levels (Figure 2A, panel 3) indicating that there is a DNA damage induced proliferation check in the diploid cells preventing their entry into meiosis. Expression of Cyclin E involved in G_1_-S transition is depleted in the Group D patients, further supporting this conclusion. In addition, in human testicular cancer CDKN1A is also present in the cytoplasm where it acts as an anti-apoptotic factor ^42^. Thus, higher expression of CDKN1A can potentially cause cell-cycle arrest while preventing apoptosis of the arrested spermatogonia and consequently impeding their entry into meiosis.

KLF4 is generally associated with the post-mitotic differentiation of the epithelial cells in both humans and mice ^43-46^. In this study, its expression was found to be much higher in the Group D patients. In humans, KLF4 is induced in response to cellular stress and up-regulates the transcription of CDKN1A and subsequently arrests cell cycle by repressing the cyclins ^47^. The over-expression of KLF4 (predominantly expressed in the non-dividing cells ^48^) in the actively dividing cells results in cell-cycle arrest with corresponding up-regulation of CDKN1A. A similar occurrence is seen in the azoospermic men with a pre-meiotic arrest (Figure 2A). KLF4 action mediated by CDKN1A induces proliferation arrest, while lack of CDKN1A results in tumorigenesis ^23,48^. Overexpression of both these genes, as seen in Group D patients, probably results in arrest of spermatogonial proliferation and differentiation halting the entire differentiation at the spermatogonial stage. This induction of CDKN1A is independent of p53 which corroborates our observation that there was no change in p53 transcript levels across the different patient groups.

Another up-regulated gene in the Group D with a potential role in spermatogenic arrest was GADD45A. In mice, this gene is expressed in the primordial germ cells (PGC) during cell fate determination, causing cell-cycle arrest ^49^. The PGCs arrested at the G2-M phase migrate to the gonadal ridge where subsequent down-regulation of GADD45A allows further proliferation confirming its negative correlation to cell proliferation ^50^.

GADD45A is also involved in the murine adult stem cell differentiation ^51^ and is up-regulated in the cells exposed to genotoxic stress causing cell-cycle arrest ^24^. Thus, the higher level of GADD45A in Group D (Figure 1A) is probably one of the factors responsible for spermatogonial cell-cycle arrest. Correspondingly, the levels of Cyclin-B are also depleted in these patients. Further, GADD45A-null cells show higher sensitivity to apoptosis under conditions of stress^52^ indicating its role as an anti-apoptotic factor and supports lack of apoptosis seen in the diploid arrested patients (Figure 3A).

There was a very significant increase in FOS and JUN expression in the Group D patients. Such an increase in FOS expression causes loss of mouse germ cells ^25^ with decreased number of spermatocytes and spermatids ^53^ which correlates with the decreased testicular cell numbers as observed in this study.

The network of diploid-enriched genes provides a list of genes that act in tandem to regulate the diploid cell population and alteration in their expression halts diploid to double-diploid transition (Figure. S9A). From our observations, both cell-cycle check points (G1-S: CDKN1A, KLF4 and G2-M: GADD45A) appear to be operational in patients with a pre-meiotic arrest. Further, the altered expression of KLF4, CDKN1A, GADD45A and FOS (important regulators of mitotic cell-cycle) could potentially be responsible for the arrest. Besides the genes validated in this study, microarray analysis and subsequent network mapping revealed that the components of the cell-cycle such as CDC25B ^26^ and FOXM1 ^54^ (crucial for expression of cell-cycle genes in G2 phase; depletion causes cell-cycle arrest) were also down regulated in the arrested patients. CDKN1A and FOXM1 are known to act in opposition to each other with CDKN1A overexpression leading to low levels of FOXM1 and disruption of the normal cell-cycle^27^. Taken together, these genes represent the potential causes of the pre-meiotic arrest.

The double-diploid-enriched genes identified in the study showed enrichment for regulation of the inflammatory response and several related pathways such as toll like receptor, TNF and MAPK signaling (Tables S3 and S4). The testes are primarily immune-privileged organs and any inflammation altering this immune-suppressed microenvironment leads to disruption of spermatogenesis ^28^. Secretion of inflammatory cytokines by circulating macrophages invading the testis in orchitis, affect the integrity of the Sertoli cell tight junctions and facilitate shedding of the immature germ cells into the lumen of the tubule causing infertility ^55-57^. CCL3 is known to attract and induce extravasation of immune cells within the testis ^58^ and its elevated levels (Figure 1B) directly correlates with the accumulation of immature double-diploid cells in the tubular lumen of Group DT patients. Among the interleukins, IL1A is normally expressed in the testis regulating the function of the blood-testis barrier ^59^ while the inflammatory cytokines show minimal expression ^29^. In rats, infertility induced by ionizing radiations (IR) in mediated by IL1B (not expressed in the normal rat testis). Further, IL1B injection alone was able to mimic the effects of IR mediated loss of fertility ^60^. IL1B also showed increased expression in the Group DT patients indicating that there is an acute inflammation in these patients (Figure 1B). Thus, the observed overexpression of the pro-inflammatory genes such as CCL3 (chemokine ligand 3), IL1B and IL8 (interleukins), could potentially alter the micro-environment within the testis and be responsible for the meiotic arrest of spermatocytes in the Group DT patients.

EGR2, an early gene expressed in the Spermatogonium-A, is known to be responsible for maintenance of spermatogonia ^61^ by controlling mitosis ^62^ and its expression must be down regulated at the onset of meiosis ^30^. As seen in the Figure 2B, expression of EGR2 persists in the nucleus of the spermatocytes in Group DT where its continued expression might provide conflicting differentiation cues to the cell resulting in disruption of meiosis and subsequent arrest. Additionally, the transcripts of FOXM1, JUND (genes involved in mitotic cell-cycle control), comprising the double-diploid network, show no change in the Group DT as compared to the Group DTH. This further validates the classification of genes as enriched in a particular testicular cell population.

Expression of majority of the haploid-enriched genes was not seen in the Groups D and DT (Figure. S9C) and can be attributed to the absence of haploid cells. To understand the role of the haploid-enriched genes in the context of spermatocyte differentiation, only those genes reported to be expressed in spermatocytes and persisting till spermatid stage were selected from the haploid-enriched genes for further analyses. These selected genes showed enrichment for pathways involved in sperm energy-metabolism and architecture (Table S4) confirming the strategy for population-enriched (D, T and H) classification employed.

One of the most interesting genes identified in this study is RFX2, a member of the RFX family of transcription factors. Expression of RFX2 is testis-specific, particularly in the meiotic and post-meiotic cells ^63^ and controls transcription of various pachytene spermatocyte-specific genes including histoneH1 ^31^. RFX2 binds to MYBL1 and broadens its action in the control of meiosis ^64^ and MYBL1 knockout mice exhibit arrested spermatogenesis at the pachytene stage ^53^. Expression of RFX2 is down-regulated in both Groups DT and D (Figure 1C). As shown in Figure 2C, expression of RFX2 was seen in the double-diploid cells of Group DTH, but not that of the Group DT suggesting its crucial role in the progression of meiosis. The complete absence of any double-diploid cells in Group D corresponds to the observed lack of staining. The role of RFX2 as the master regulator of spermiogenesis ^65^, controlling the genes required for ciliogenesis ^66^ is well established. However, its role in the control of meiosis is less studied and requires investigation. The present study suggests a greater role for RFX2 in regulating the double-diploid-haploid transition which is in agreement with Kistler *et al* who suggested a similar role for RFX2 in rats^64^.

CST8 belonging to the cystatin superfamily of cysteine protease-inhibitors is mainly expressed in the testis and epididymis^67^. CST8-null mice show a defect in the Sertoligerm-cell adhesion and premature release of germ cells into the lumen with the severity increasing with age^32^. In the histological sections of the DT patients, a large number of the double-diploid cells were present in the lumen of the tubules with a disruption of proper tubular architecture similar to that seen in CST8-null mice (unpublished data) indicating a similar role for CST8 in human spermatogenesis. Lack of expression of CST8 (Figure 1C) might be responsible for loss of germ-cell integrity, disruption of the Sertoli cell-germ cell interactions and shedding of spermatocytes in the tubular lumen of the Group DT patients.

The alterations in expression of genes such as DDIT3, ATFs, GADD45A, and CDKN1A in the Group DT patients strongly suggest that DNA damage is extant in the germ cells. In this context, the loss of GGN (gametogenetin) in the Group DT (Figure 1C) indicates its probable contribution towards the genotoxic stress. GGN, a mouse pachytene spermatocyte specific gene, along with POG (proliferation of germ cells), controls meiosis by regulating the DNA double strand break repair^68^. GGN-null mice are embryonic lethal while GGN^+/−^ mice are infertile with an arrest at the pachytene stage as the DNA double strand breaks during homologous recombination are not repaired in these mice^33^. The Group DT patients show an arrest in spermatogenesis at the meiotic stage and a down-regulation of GGN at the transcript level indicating that the hampered DNA damage repair and induction of DNA damage inducible cell-cycle arrest together cause the meiotic arrest.

Intriguingly, the androgen receptor gene, which is expressed in the Sertoli cells, was observed to be differentially expressed across all the networks indicating that disruption of germ-cell development has a direct bearing on the Sertoli cells.

As seen in Figure 3, autophagy was up-regulated in the Groups D and DT patients as compared to the Group DTH. This observation indicated that autophagy was a probable mechanism for clearance of the arrested cells in NOA and the regulation seems to be at the protein level. Interestingly, CDKN1A is reported to induce autophagy^69^ and increased autophagy could be a consequence of its observed high levels. Our data suggests a role for autophagy in NOA and male infertility which requires further investigation.

This study suggests that the major disruption in azoospermia is in the transition from one phase to another. In Group D patients, the arrest seems to be a result of alterations of genes such as CDKN1A, KLF4, GADD45A and FOS regulating mitosis, consequently, the diploid to double-diploid transition. In Group DT patients, alterations of genes such as RFX2, GGN and CST8 that were down-regulated and persistence of EGR2 could be the potential causes of meiotic arrest. The present study suggests that the regulation of the diploid - double-diploid - haploid transition are multigenic with the tandem alteration of several genes resulting in infertility (Figure 4). The exact point of arrest in humans within a phase still remains to be identified.

**Figure 4.**
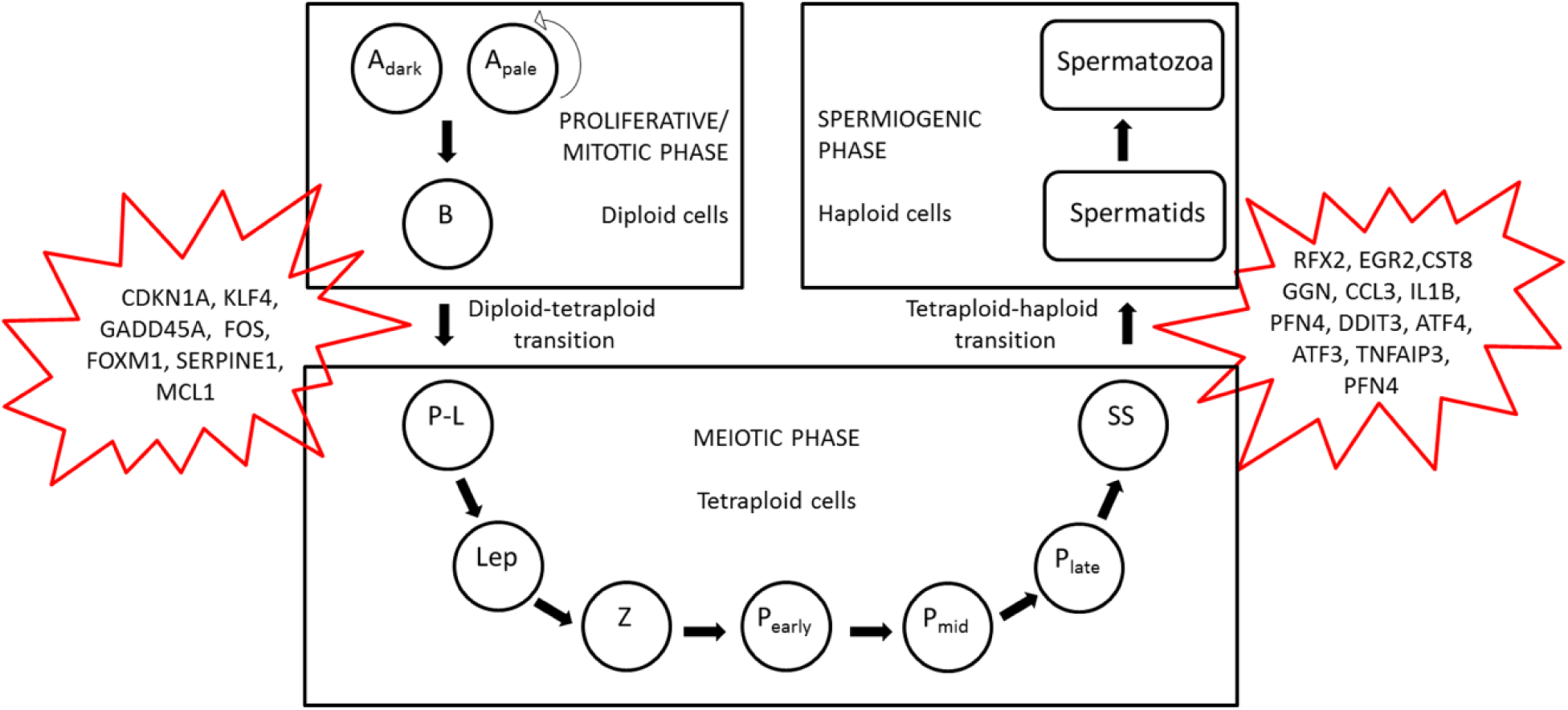
Schematic representation of the progression of human spermatogenesis and its relationship with the altered population-enriched genes identified from this study. The phases of human spermatogenesis as well the points of transition between each phase are shown along with the altered genes identified from this study as potential causes of the spermatogenic arrest. Adark: dark A type Spermatogonia, Apale: pale A type Spermatogonia capable of self-renewal marked by arrow, B: Type B spermatogonia, P-L: pre-leptotene spermatocytes, Lep: leptotene spermatocytes, Z: zygotene spermatocytes, Pearly: early-pachytene spermatocytes, Pmid: mid-pachytene spermatocytes, Plate: late-pachytene spermatocytes, SS-secondary spermatocytes.

In conclusion, this study provides glimpses into understanding of the possible mechanisms involved in the regulation of human spermatogenesis and their relation to infertility. The genes that have been analyzed using both human and rodent data in this study are restricted to those with most striking changes in expression. There is a possibility that the genes with relatively smaller differences in expression levels may have profound effects on spermatogenesis and hence need to be analyzed further. The microarray analysis associated with this study provides a huge resource for analyzing such genes and further analysis may lead to identification of other potential targets for infertility treatment. More detailed analysis of genes responsible for arrest of spermatogenesis can also lead to identification of potential targets for male contraception.

## Additional Information

### Author Contributions

The research described here was designed and executed by AB and RRD. SSV identified the infertile patients and provided the biopsies. RM was involved in autophagy experiments. RJ and PK analyzed the microarray data and RRD supervised the entire project. AB, PK and RRD prepared the manuscript. All authors read and approved the manuscript.

## Acknowledgements

The authors thank Prof. M.R.S. Rao, JNCASR for scientific discussions and Prof. N.V. Joshi, CES, IISc for help with the statistics.

---

**Supplementary material for manuscript - ‘Correlation between the aberrant human testicular germ-cell gene expression and disruption of spermatogenesis leading to male infertility’**

**List of Primers used for real time PCR**

**Table S1.**
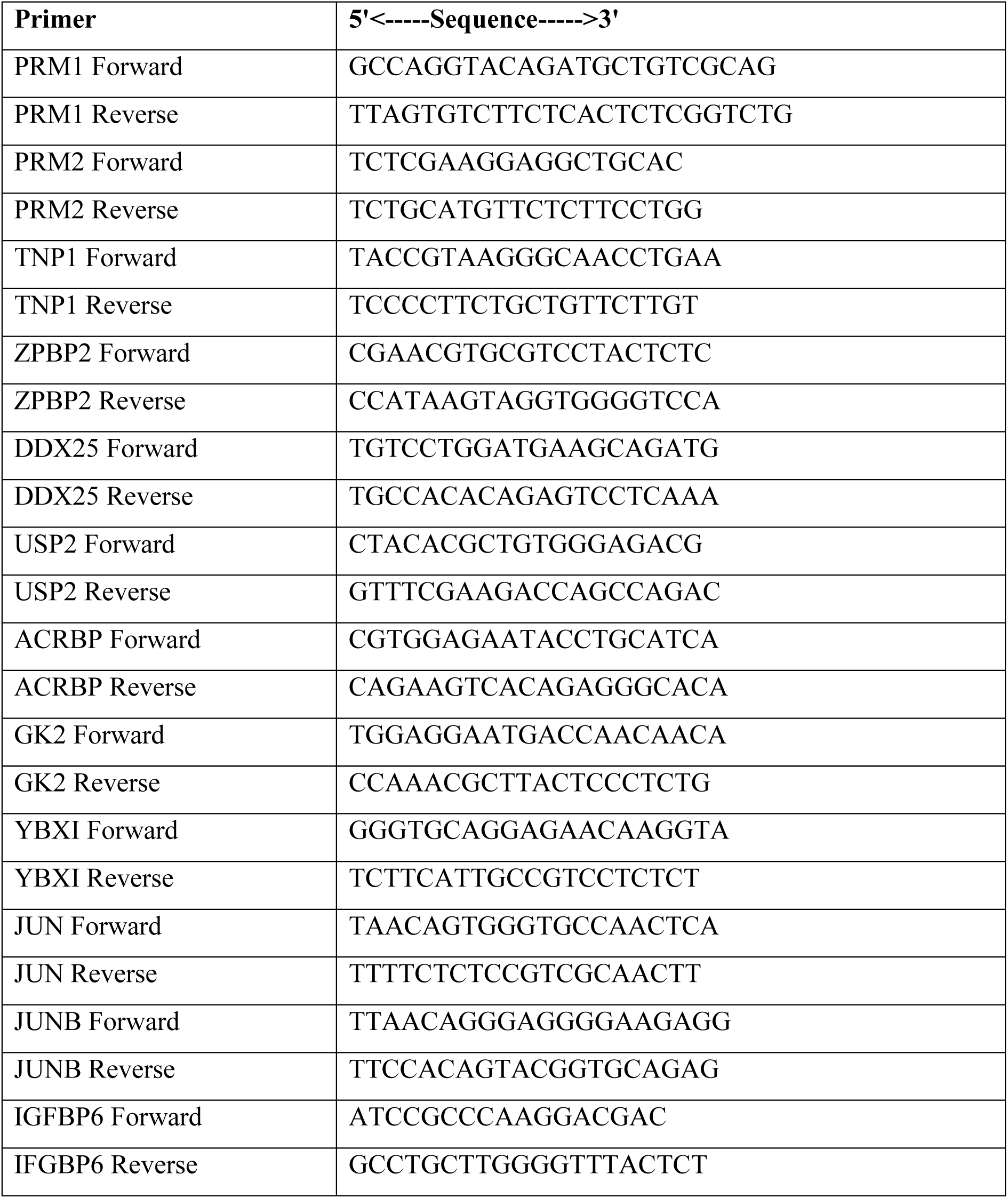
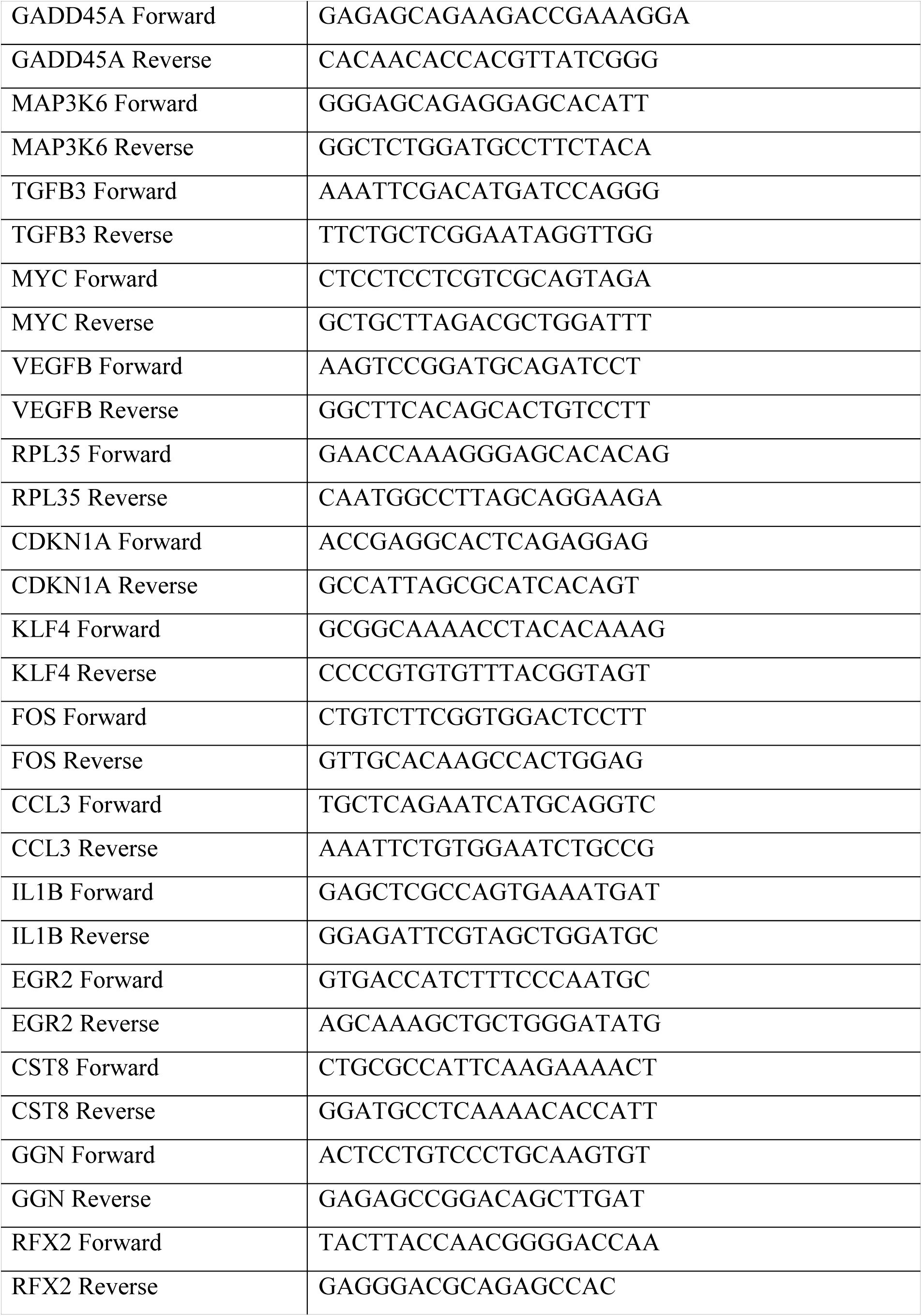
(Supplementary Table 1)

## Extended Materials and Methods

### Preprocessing and Identification of differentially-expressed genes

Average signal intensities of all the genes were background-subtracted and Quantile normalization was applied between the arrays. The Linear model and empirical Bayes methods were performed to assess the differentially-expressed genes between the Normal and the patients’ samples with arrested spermatogenesis using ‘Limma’ (Linear Models for microarray data) package in R-3.3.2 (1). The Benjamini and Hochberg method was used to correct for multiple testing(2) within Limma. The genes with adjusted P-values <0.05 and fold change >1.5 were considered significantly regulated. All analyses were represented as ‘Y’VS‘X’ where X is the arrested population (variable) and Y is the normal population (constant). (Table S1)

The differentially-expressed genes with fold changes >3 from the above analysis were further analyzed using ‘Venny’ (bioinfogp.cnb.csic.es/tools/venny/index.html)(3) and their individual fold changes (from Microarray) were plotted manually for identifying the genes commonly and differentially-expressed across all sample groups.

## Immunohistochemistry

Paraffin sections (4 μm) were embedded on silane-coated slides for Immunohistochemistry. Paraffin was removed and sections were rehydrated by passing through xylene (15 minutes), decreasing concentrations of ethanol (100%, 95%, and 70%; 5 minutes each) and double distilled water (5 minutes). Antigen-retrieval was carried out in citrate buffer (pH-6) by heating the sections in a microwave oven at 600 W for 20 minutes. The sections were cooled to room temperature, washed in 1X PBS and the intracellular peroxidase was quenched by methanol-H_2_O_2_ treatment for 20 minutes at room temperature. The sections were washed again with PBS and non-specific binding was blocked by 5% skim milk and incubated with the primary antibodies (at recommended dilutions) overnight at 4 °C. The signal was developed using the PathSitu PolyExcel HRP/Dab Detection System kit. The sections were subsequently dehydrated and mounted in DPX. Sections incubated with only secondary antibody served as negative controls.

## TUNEL assay

The slides were rehydrated in different gradients of alcohol and finally immersed in TBS. The sections were permeabilized using a Proteinase K solution followed by the quenching of the endogenous peroxidases by H_2_O_2_-methanol. The sections were labeled with a TdT labeling mix in a humidified chamber for 90 minutes. The reaction was stopped by adding the Stop buffer and the samples were washed in TBS. Streptavidin-HRP was added and the reaction developed using Dab solution. The slides were washed, air-dried and mounted in DPX post development. DNase treated section served as positive control whereas TdT untreated sections were served as negative control.

**Table S2.**
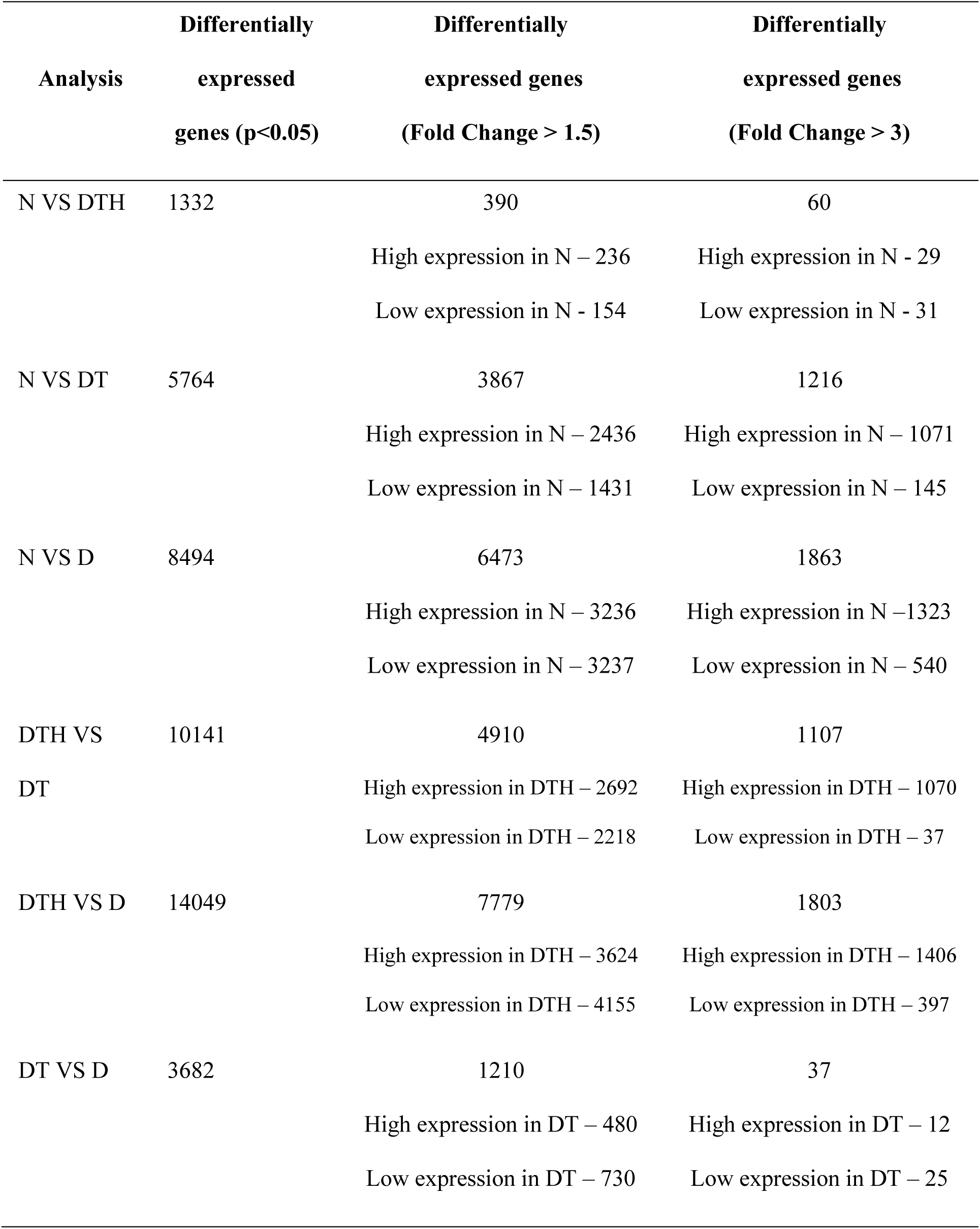
(Supplementary Table 2) N: Pooled RNA control, DTH: obstructive azoospermia patients with complete intra-testicular spermatogenesis, DT: meiotic arrest patients with diploid and double-diploid cells in their testes, D: pre-meiotic arrest patients with only diploid cells in their testes.

**Table S3.** (Supplementary Table 3) Gene enrichment analysis for the cell population specific genes: The different cell population enriched gene sets were analyzed for enrichment to GO terms to verify whether the classification as diploid, double-diploid and haploid specific genes was accurate

| DIPLOID-ENRICHED |  | DOUBLE-DIPLOID-ENRICHED |  | HAPLOID-ENRICHED |  |
| --- | --- | --- | --- | --- | --- |
| Biological process | p value | Biological Process | p value | Biological process | p value |
| response to steroid hormone | 1.02E-06 | regulation of inflammatory response | 4.23E-05 | sperm motility | 6.7E-09 |
| response to corticosteroid | 1.32E-05 | negative regulation of apoptotic signaling pathway | 0.000217 | cilium movement | 2.08E-05 |
| regulation of cell growth | 2.48E-05 | MAPK cascade | 0.00015 | male gamete generation | 1.39E-05 |
| negative regulation of extrinsic apoptotic signaling pathway | 0.000292 | cellular response to gonadotropin stimulus | 0.000373 | spermatogenesis | 1.31E-05 |
| response to progesterone | 0.000245 | regulation of cellular response to stress | 0.000421 | spermatid development | 1.49E-05 |
| transcription from RNA polymerase II promoter | 0.001754 | cellular process involved in reproduction | 0.00046 | gamete generation | 9.14E-05 |
| germ cell migration | 0.000727 | positive regulation of cell cycle | 0.00096 | sperm-egg recognition | 1.53E-05 |
| cell cycle arrest | 0.015158 | positive regulation of cellular component movement | 0.001219 | axonemal dynein complex assembly | 0.001794 |

**Table S4.** (Supplementary Table 4) Enriched pathways for the cell population enriched genes: Pathway analysis through KEGG provided the pathways regulated by the population enriched genes. In case of the diploid-enriched genes, the major associated pathways included growth development and the double-diploid-enriched genes showed enrichment for the meiotic, inflammatory and apoptotic pathways. The genes that showed enrichment for pathways such as spermatid maturation, sperm formation and architecture, ciliary and flagellar organization as well as energy production all belonged to the haploid-enriched gene set

| **DIPLOID-ENRICHED** | **DOUBLE-DIPLOID-ENRICHED** | **HAPLOID-ENRICHED** |
| --- | --- | --- |
| **Biological Process** | **Biological Process** | **Biological process** |
| TNF signaling pathway | Toll-like receptor signaling pathway | Glycolysis / Gluconeogenesis |
| MAPK signaling pathway | Rheumatoid arthritis | Pentose phosphate pathway |
| p53 signaling pathway | TNF signaling pathway | Biosynthesis of amino acids |
| Cell cycle | MAPK signaling pathway | Carbon metabolism |
| PI3K-Akt signaling pathway | Graft-versus-host disease | Tight junction |
| Oxytocin signaling pathway | Cytokine-cytokine receptor interaction | Metabolic pathways |
| GnRH signaling pathway | Oocyte meiosis | Endocytosis |
| NF-kappa B signaling pathway | NOD-like receptor signaling pathway | Starch and sucrose metabolism |
| Estrogen signaling pathway | Protein processing in endoplasmic reticulum | Regulation of actin cytoskeleton |
| VEGF signaling pathway | Chemokine signaling pathway | Fanconi anemia pathway |
| Jak-STAT signaling pathway | Estrogen signaling pathway | Fatty acid biosynthesis |

**Table S5.** (Supplementary Table 5) Status of Apoptotic pathway genes across transcriptome analyses

| Gene | N VS DTH | N VS DT | N VS D | DTH VS DT | DTH VS D | DT VS D |
| --- | --- | --- | --- | --- | --- | --- |
| BCL2 | NC | -0.89 | -0.76 | NC | -0.88 | -0.73 |
| BCL2L1 | NC | NC | NC | NC | NC | NC |
| BAX | NC | NC | NC | NC | -0.76 | NC |
| BAD | NC | NC | NC | NC | NC | NC |
| BID | NC | NC | NC | NC | NC | NC |
| APAF1 | NC | NC | NC | NC | NC | NC |
| CASP8 | NC | NC | NC | NC | NC | NC |
| CASP9 | NC | NC | NC | NC | NC | NC |
| CASP3 | NC | NC | NC | NC | NC | NC |
| CASP6 | NC | NC | NC | NC | -0.59 | NC |
| CASP7 | NC | NC | NC | NC | NC | NC |
| FAS | NC | NC | -1.06 | -0.63 | -1.20 | NC |
| FASLG | NC | NC | NC | NC | NC | NC |

*Note:* NC- no change in expression; Fold change (FC) is represented as $\log_2$ FC; Differential expression is from samples analyzed in microarray.

**Table S6.** (Supplementary Table 6) Status of Autophagy genes across transcriptome analyses

| **Gene** | **N VS** | **N VS DT** | **N VS D** | **DTH VS** | **DTH VS D** | **DT VS D** |
| --- | --- | --- | --- | --- | --- | --- |
|  | **DTH** |  |  | **DT** |  |  |
| ATG3 | NC | 1.44 | 0.86 | NC | 1.08 | NC |
| ATG4C | NC | 1.03 | NC | NC | NC | NC |
| ATG16L2 | NC | -0.73 | NC | NC | -0.67 | NC |
| MAP1LC3A | NC | NC | NC | NC | NC | NC |
| ATG2A | NC | -0.97 | -0.86 | NC | NC | NC |
| ATG9A | NC | NC | NC | 0.96 | 0.83 | NC |
| ATG 7 | NC | NC | NC | 0.61 | NC | NC |
| DRAM1 | NC | -1.14 | NC | -0.97 | -1.43 | NC |
| GABARAPL1 | NC | -1.14 | NC | -0.85 | NC | NC |
| BECN1 | NC | NC | NC | NC | -0.59 | NC |
| PIK3R2 | NC | -0.85 | -0.84 | NC | NC | NC |
| PIK3AP1 | NC | NC | NC | NC | -0.63 | NC |
| PIK3R1 | NC | NC | NC | NC | -0.81 | NC |

*Note:* NC- no change in expression; Fold change (FC) is represented as log<sub>2</sub> FC; Differential expression is from samples analyzed in microarray.

**Figure S1.**
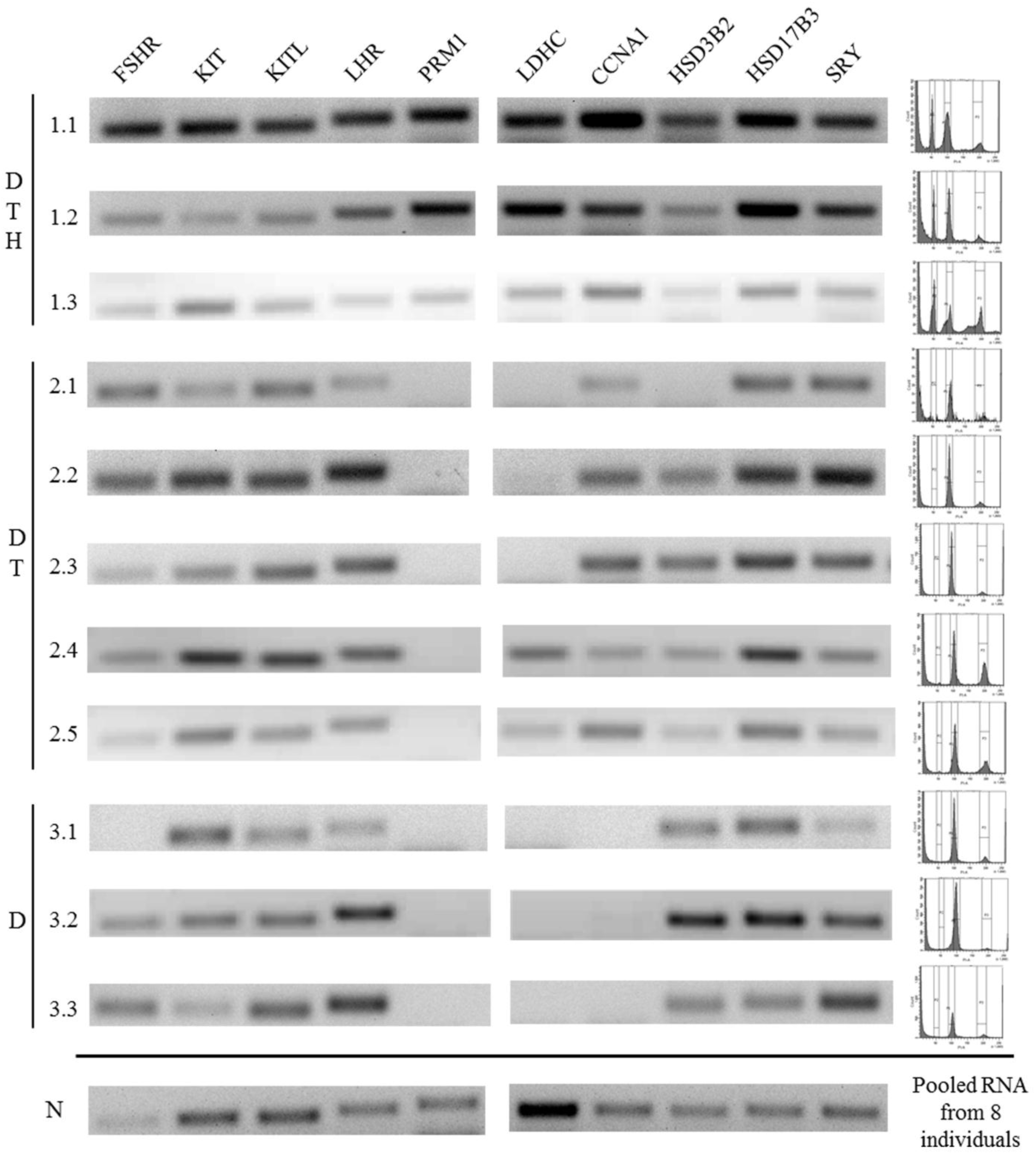
(Supplementary Figure 1) Legend: Germ cell patterns of patient samples used for transcriptome analysis ascertained by flow cytometry and amplification of marker genes. DTH: obstructive azoospermia patients with complete intra-testicular spermatogenesis, DT: meiotic arrest patients with diploid and double-diploid cells in their testes, D: pre-meiotic arrest patients with only diploid cells in their testes, N: Pooled RNA control.

**Figure S2.**
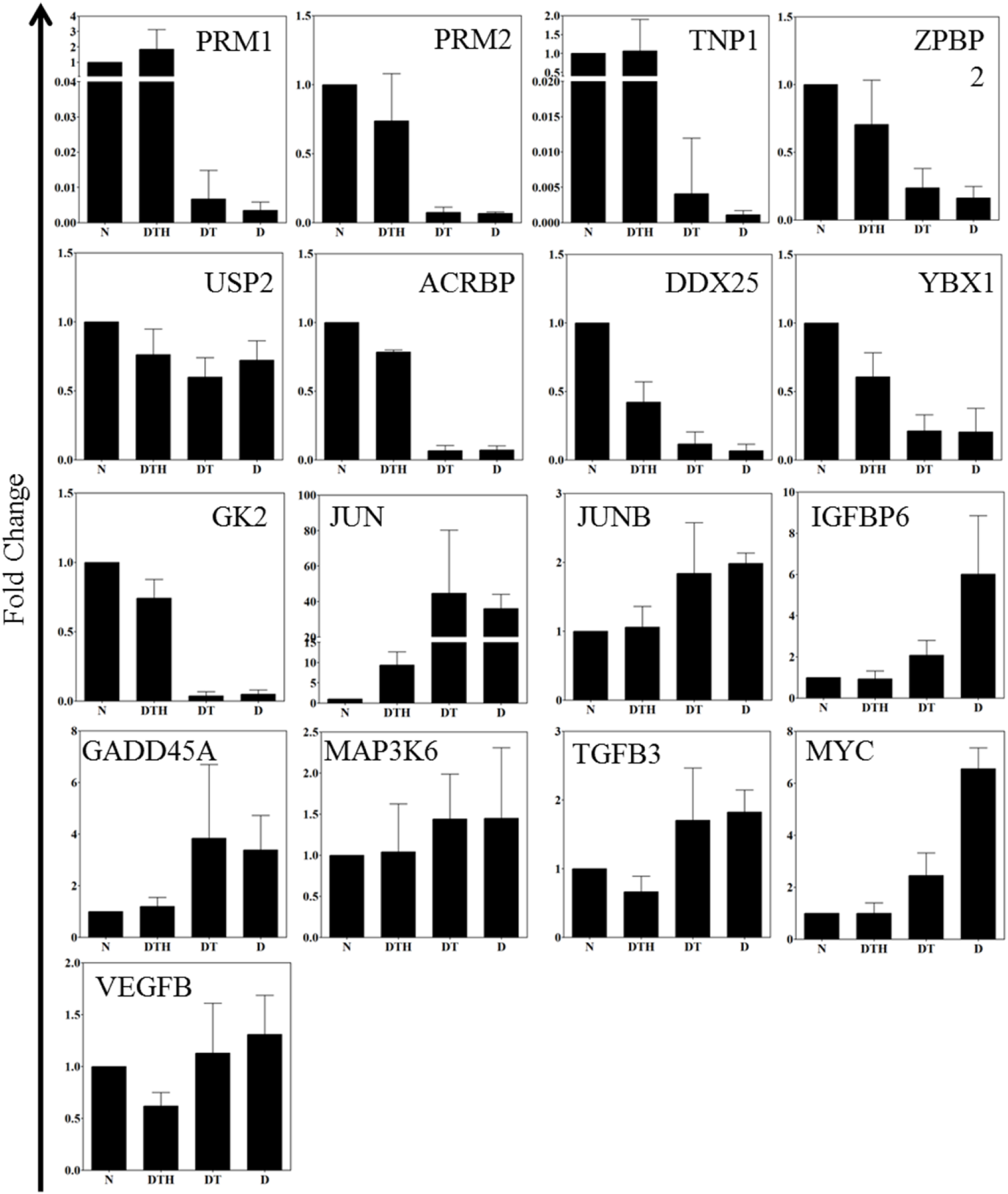
(Supplementary Figure 2) Legend: Validation of microarray. A total of top 17 differentially expressed genes, identified from the transcriptome analysis (D Vs N and DT Vs N) that also are the known markers of individual testicular cell populations, were quantitated in qRT-PCR. The Y-axis represents the fold changes as compared to N (which is plotted as 1). PRM1 and PRM2 and TNP1 (4-6), the known markers for the haploid population were not expressed in samples from Group DT and D. ZPBP2, ACRBP, USP2 involved in sperm motility, DDX25 essential for spermatid development, GK2 energy pathway, and YBX1 (7-9), known to be expressed in the mature spermatids, were also expressed more in N and DTH as compared to the groups with a spermatogenic arrest. Important transcription factors such as FOS, JUN and growth factors such TGFB3 and IGFBP6 (involved in growth inhibitory signal from the Sertoli Cell) mostly were expressed more in DT and D as compared to DTH and N. MYC, a marker for spermatogonial cells (10) was expressed more in the testes of patients belonging to Group DT and D. Also there was no significant difference in the expression levels of the validated genes between N and patient samples belonging to Group DTH. However, for further analysis N and Group DTH were maintained as separate entities and all comparisons of the arrested samples (Group DT and D) were made with respect to N. [N: Pooled RNA control, DTH: obstructive azoospermia patients with complete intra-testicular spermatogenesis, DT: meiotic arrest patients with diploid and double-diploid cells in their testes, D: pre-meiotic arrest patients with only diploid cells in their testes.]

**Figure S3.**
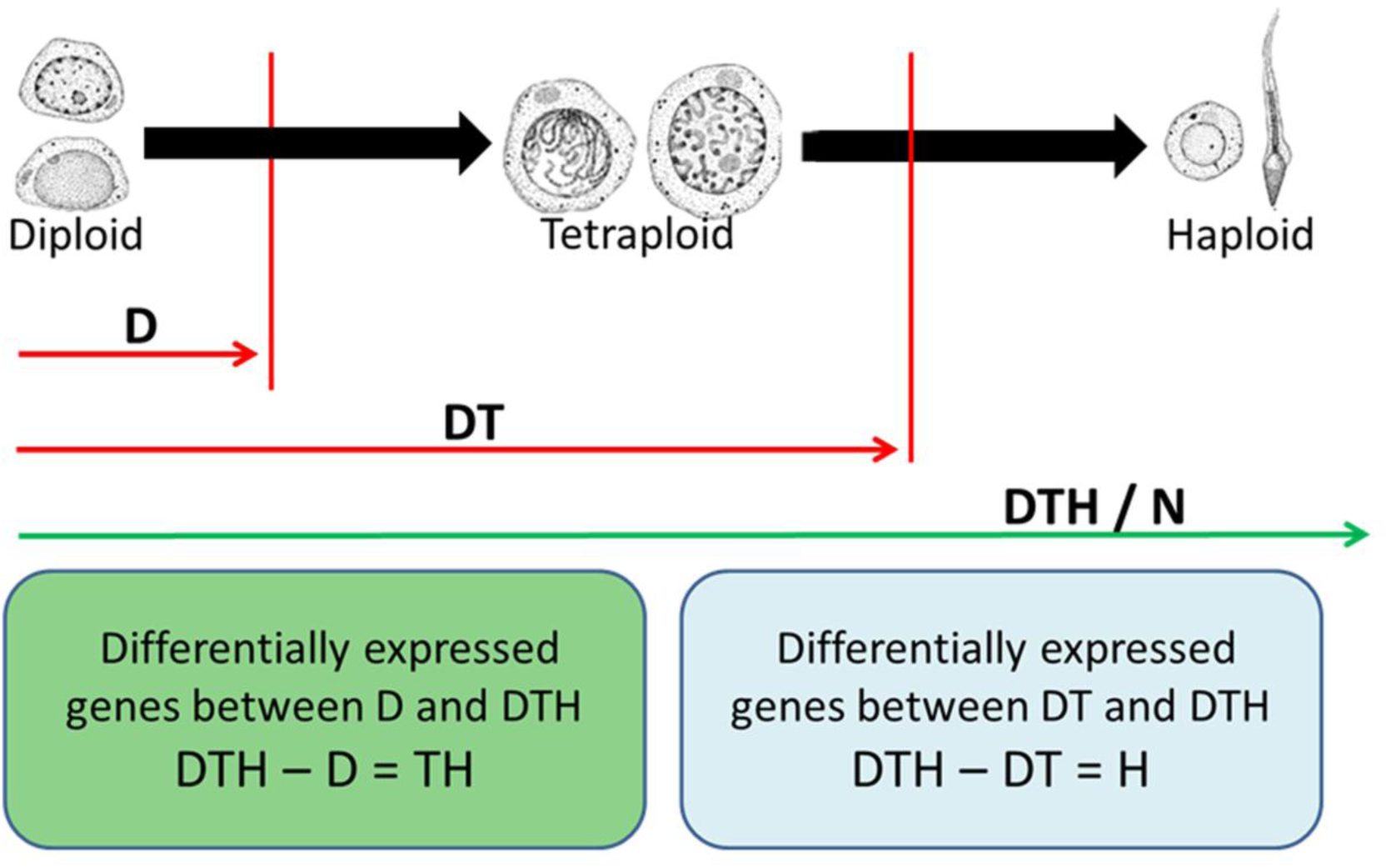
(Supplementary Figure 3) Legend: Schematic of microarray data analyses. The Double-diploid and haploid-enriched genes were arrived at by subtracting the differentially expressed genes in the Groups D and DT from the differentially expressed genes in the control pooled RNA (N). The differential gene expression between N (cell populations present: DTH) and Group D (cell populations present: D) correspond to the double-diploid and haploid-enriched genes (DTH-D=TH) [N VS D], while the genes differentially expressed between N and Group DT (cell populations present: DT) represent the haploid population–enriched genes (DTH-DT=H) [N VS DT]. Further, the comparison of these two differentially expressed gene sets was carried out to identify the double-diploid-enriched genes (the common genes between N VS D and N VS DT represent the set of haploid-enriched genes and the remainder would be double-diploid-enriched genes). D: diploid cells; T: double-diploid cells, H: haploid; Group D: pre-meiotic arrest patients with only diploid cells in their testes; Group DT: meiotic arrest patients with diploid and double-diploid cells in their testes; Group DTH: obstructive azoospermia patients with complete intra-testicular spermatogenesis; N: Pooled RNA control

**Figure S4.**
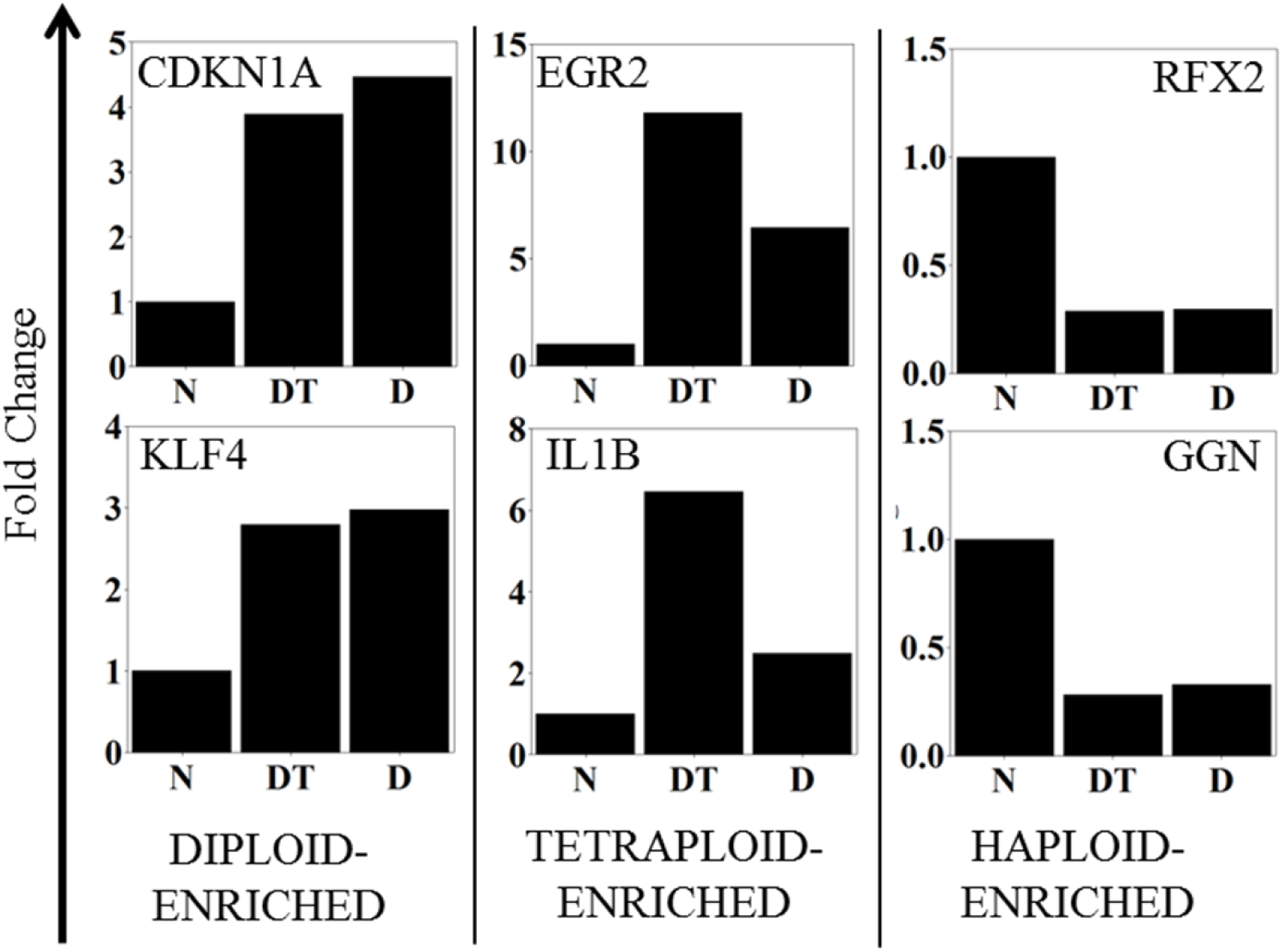
(Supplementary Figure 4) Legend: Representative graphs showing differential gene-expression pattern of the testicular cell population (diploid, double-diploid and haploid) enriched genes. The median fold change of all the 1754 genes (from N Vs D and N Vs DT analyses; Supplementary Table 2) were computed and the genes showing enrichment in a particular cell population (diploid, double-diploid and haploid) were annotated as shown. This was done to confirm that the differences seen in the transcriptome analysis were due to alterations in the gene expression itself and not a result of the presence of different cell populations in the different groups. The genes that showed minimal expression in both Groups DT and D patients but showed elevated expression in N and Group DTH patients were annotated as haploid-enriched genes while the genes with elevated expression in either Groups DT or D and lower expression in Group DTH and N were annotated as double-diploid-enriched or diploid-enriched respectively. A total of 74 diploid-enriched, 32 double-diploid-enriched and 794 haploid-enriched genes were identified.

**Figure S5.**
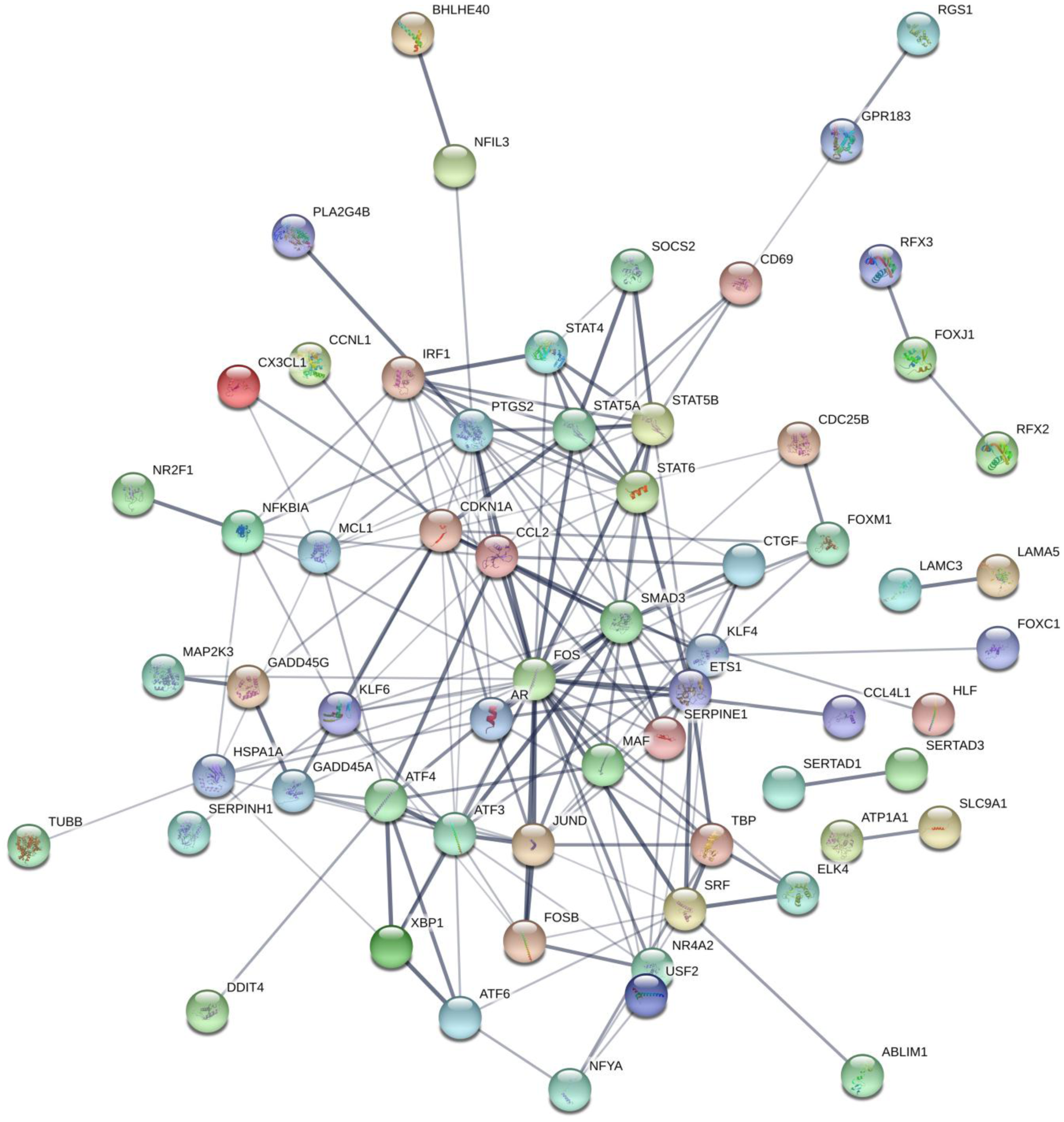
(Supplementary Figure 5) Legend: Unbiased network map of the diploid-enriched genes before selection of genes and TFs with >4 connections.

**Figure S6.**
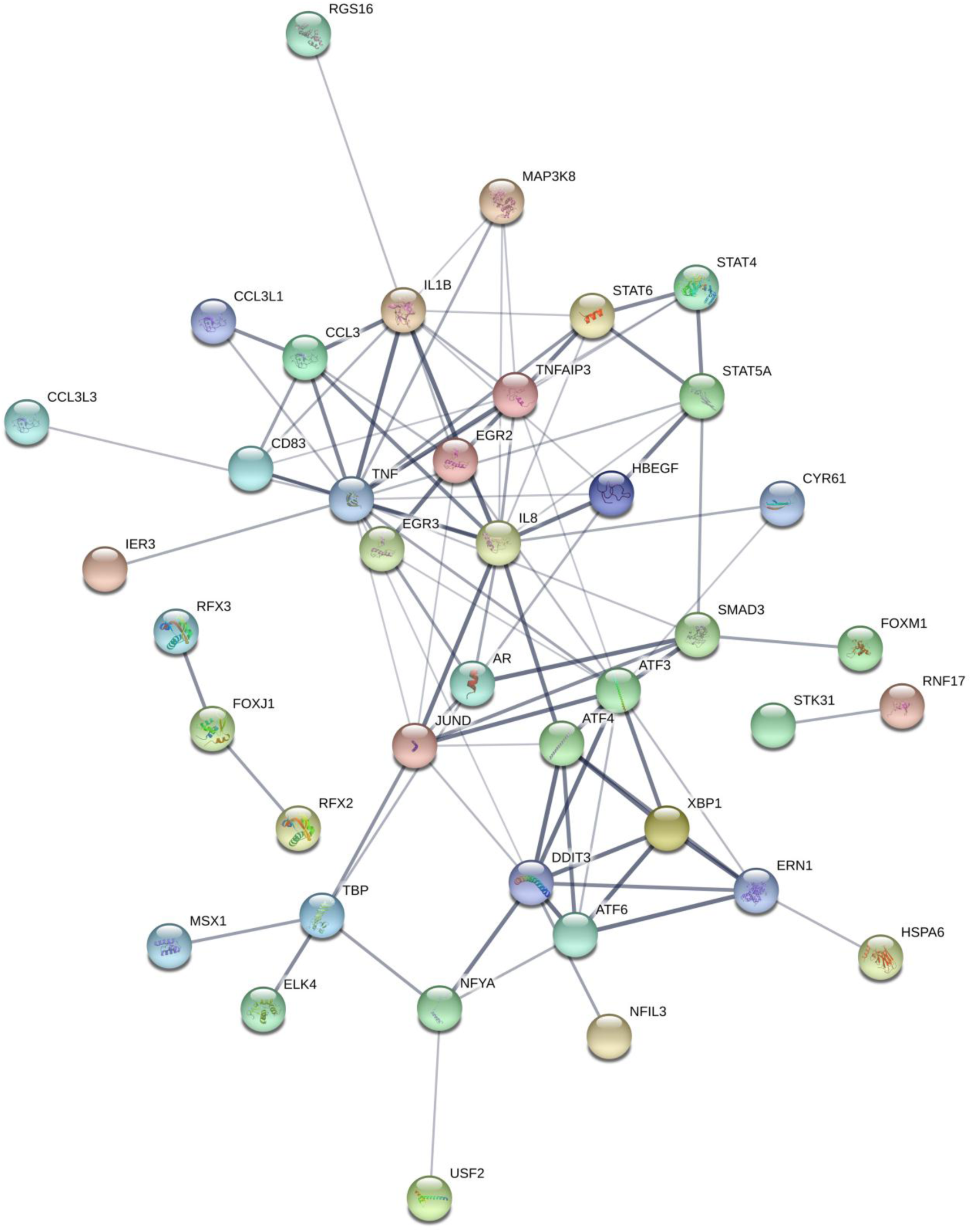
(Supplementary Figure 6) Legend: Unbiased network map of the double-diploid-enriched genes before selection of genes and TFs with >4 connections.

**Figure S7.**
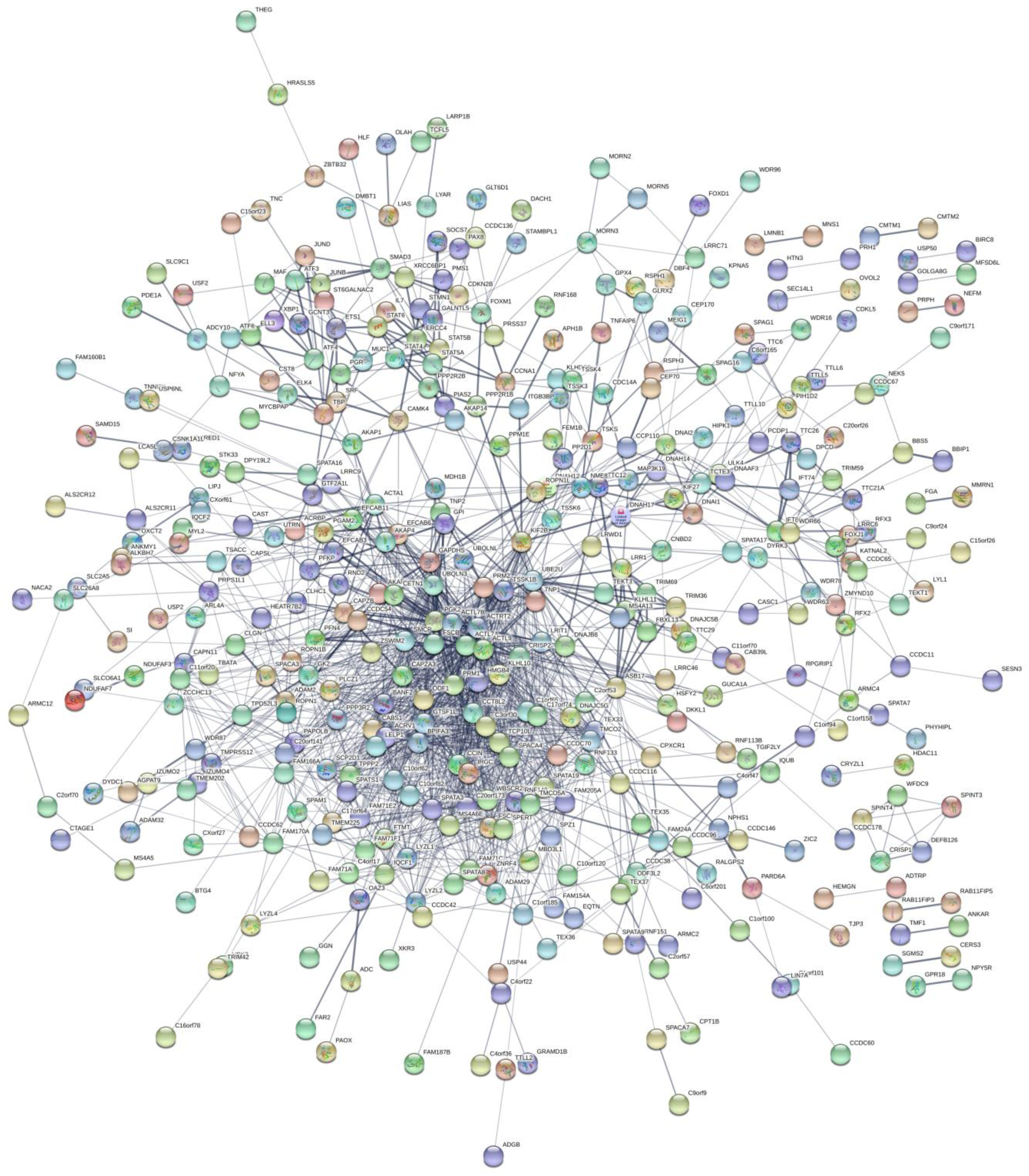
(Supplementary Figure 7) Legend: Unbiased network map of the haploid-enriched genes before selection of genes and TFs with >4 connections.

**Figure S8.**
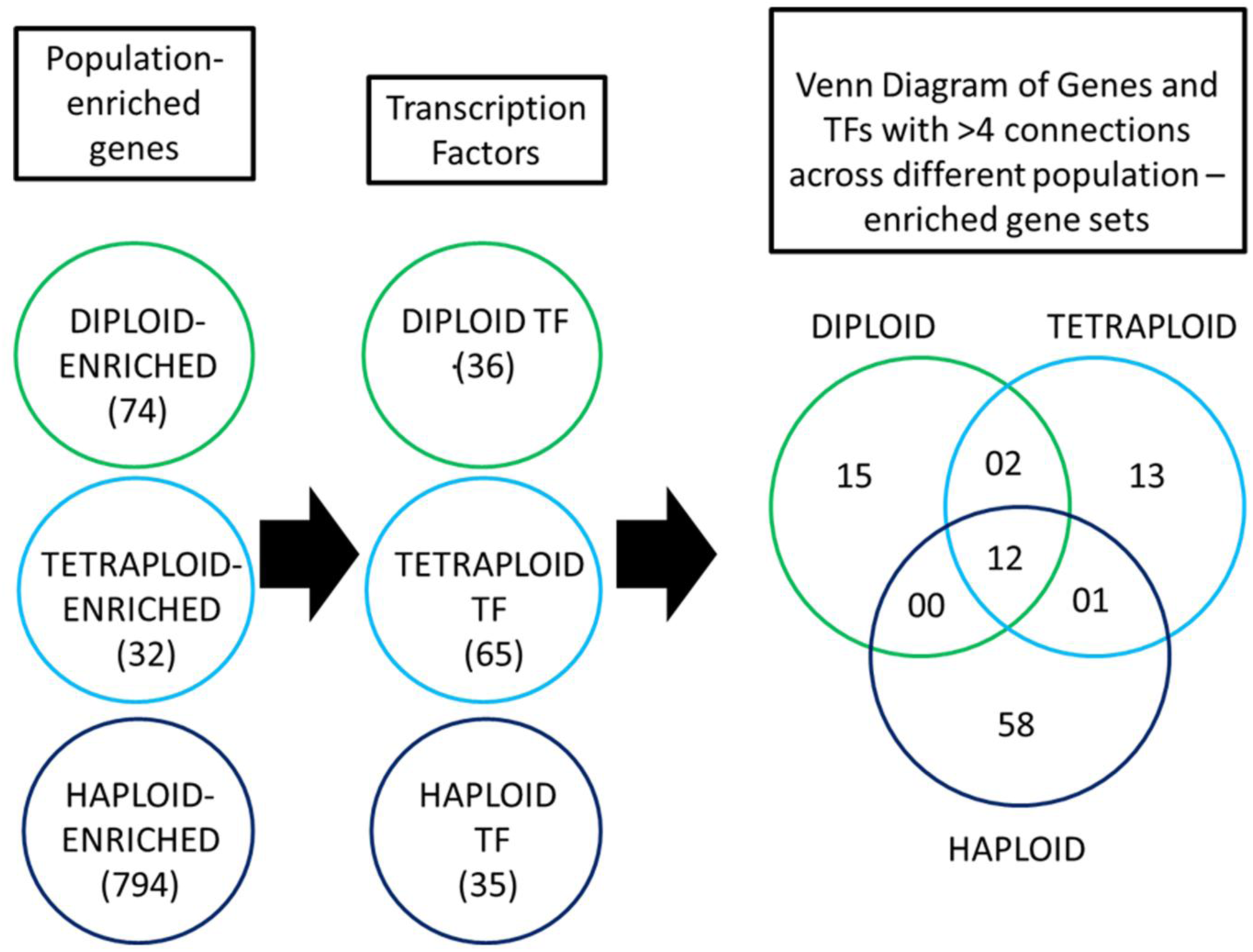
(Supplementary Figure 8) Legend: Schematic representation of workflow to identify the population-enriched genes with putative role in the spermatogenic arrest. The population-enriched gene sets were analyzed for the presence of possible transcription factor (TF) binding sites. From the putative TFs identified, the unique transcription factors differentially expressed among the different groups of patients were identified. A total of 36 diploid-enriched, 65 double-diploid-enriched and 35 haploid-enriched TF’s were identified and used to construct network maps with their respective population-enriched gene sets. The transcription factors and genes that have a large number of connections with other genes in a network represent hubs that either exert control over several genes or are the genes under tight regulation and their disruption would have wider reaching effects on the constituent pathway. Subsequent to the construction of network maps, the genes and TF’s with >4 connections were chosen. The Venn diagram represents the distribution of differentially expressed genes and TFs (both with >4 connections) across the different population-enriched gene sets. These genes represented in the Venn diagram were then further analyzed for their putative role in the spermatogenic arrest.

**Figure S9.**
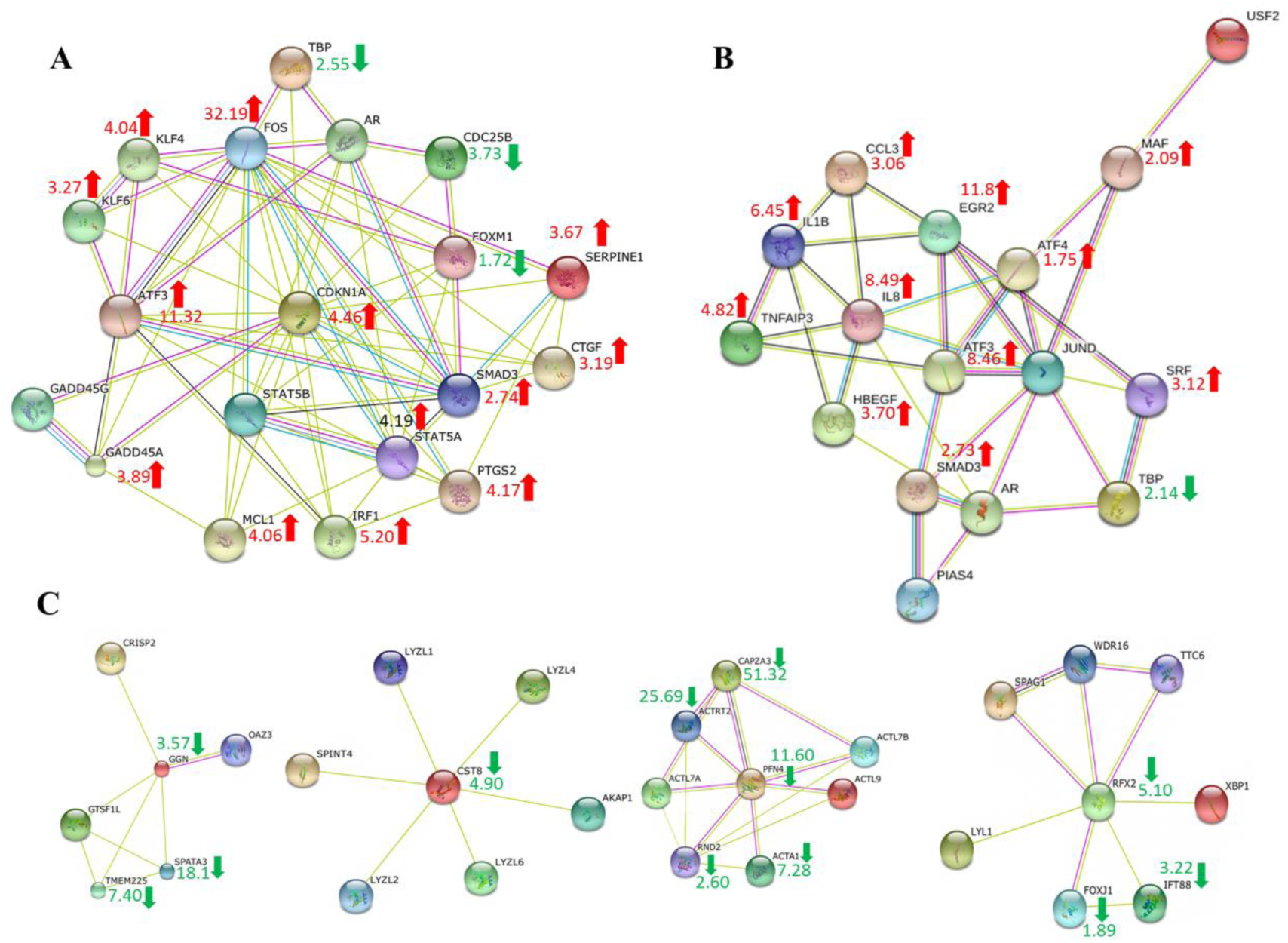
(Supplementary Figure 9) Legend: Network maps for testicular cell population-enriched genes. A. Diploid-enriched genes involved in diploid to double-diploid transition. The network map for the subset of the diploid-enriched genes involved in diploid to double-diploid transition is shown. The red arrows indicate over-expression while the green arrows indicate down-regulation in the Group D (pre-meiotic arrest patients with only diploid cells in their testes). The numerals represent the fold change of the corresponding genes (from microarray; D Vs N analysis) in Group D with respect to N (Pooled RNA control). **B. Double-diploid-enriched genes involved in double-diploid to haploid transition**. The network map for the subset of the double-diploid-enriched genes involved in double-diploid to haploid transition is shown. The red arrows indicate over-expression while the green arrows indicate down-regulation in the Group DT (meiotic arrest patients with diploid and double-diploid cells in their testes). The numerals represent the fold change of the corresponding genes (from microarray; DT Vs N analysis) in Group DT with respect to N. **C. Haploid-enriched genes involved in double-diploid to haploid transition**. The network maps for the subset of haploid-enriched genes involved in double-diploid to haploid transition is shown. The green arrows indicate down-regulation in the Group DT. The numerals represent the fold change of the corresponding genes (from microarray; DT Vs N analysis) in Group DT with respect to N.

